# Topological and Kinetic Determinants of the Modal Matrices of Dynamic Models of Metabolism

**DOI:** 10.1101/107425

**Authors:** Bin Du, Daniel C. Zielinski, Bernhard O. Palsson

## Abstract

Linear analysis of kinetic models of metabolism can help in understanding the dynamic response of metabolic networks. Central to linear analysis of these models are two key matrices: the Jacobian matrix (**J**) and its modal matrix (**M**^-1^). The modal matrix contains dynamically independent motions of the kinetic model, and it is sparse in practice. Understanding the sparsity structure of the modal matrix provides insight into metabolic network dynamics. In this study, we analyze the relationship between **J** and **M**^-1^. First, we show that diagonal dominance occurs in a substantial fraction of the rows of **J**, resulting in simple modal structures within **M**^-1^. Dominant diagonal elements in **J** approximate the eigenvalues corresponding to these simple modes, in which a single metabolite is driven back to its reference state on a characteristic timescale. Second, we analyze more complicated mode structures in **M**^-1^, in which two or more variables move in a certain ratio relative to one another on defined time scales. We show that complicated modes originate from sub-matrices of topologically connected elements of similar magnitude in **J**. Third, we describe the origin of these mode structure features based on the network stoichiometric matrix **S** and the reaction kinetic gradient matrix **G.** We demonstrate that the topologically-connected reaction sensitivities of similar magnitude in **G** play a central role in determining the mode structure. Ratios of these reaction sensitivities represent equilibrium balances of half reactions that are defined by linearization of the bilinear mass action rate laws followed by enzymatic reactions. These half-reaction equilibrium ratios are key determinants of modal structure for both simple and complicated modes. The work presented here helps to establish a foundation for understanding the dynamics of kinetic models of metabolism, which are rooted in the network structure and the kinetic properties of reactions.

## Introduction

In recent years, kinetic models of metabolism have become increasingly detailed, comprehensive, and consistent with the underlying biochemistry and genetics (1)(2)(3)(4)(5)(6). The development of kinetic models is attractive because, among other reasons, these models can help us understand the relationship between metabolite concentrations and reaction fluxes, which currently is difficult to analyze directly with constraint-based or statistical models (7)(8)(9). Kinetic models have shown utility in numerous applications, including the study of: 1) regulatory mechanisms controlling the cellular metabolic network (10)(11), 2) complex dynamic behavior such as bistability (12), 3) intracellular signal transduction (13), and 4) the effect of enzyme mutations on a network scale (14)(15). Furthermore, predictive kinetic models are desirable in metabolic engineering to improve production, substrate utilization, and product quality (16)(17).

A grand challenge for the further development of kinetic models is to develop a fundamental understanding of their behavior, in order to enable a deeper understanding of the structure and function of the biological system. A number of studies have made theoretical and practical headway in this regard by analyzing the linear properties of the dynamic system around a steady state. These linear analysis methods have helped to provide insight into metabolic flux control (18)(19), elucidate the temporal hierarchy of dynamic events (20), and determine the fundamental dynamic structure of the network (21).

At the core of these linear analysis methods is the modal matrix (**M**^-1^) resulting from the Jacobian matrix (**J**) of the mass balance equation. The modal matrix contains dynamically decoupled motions of the metabolic network, called modes. For real metabolic networks, the modal matrix has a sparse structure (20), the interpretation of which can yield biological insight into dynamics occurring on particular time scales. However, while **M**^-1^ is a numerically-generated matrix, **J** can be represented symbolically in terms of derivatives of the reaction rate laws (*d***v**/*d***x**) in the network. Thus, obtaining an understanding of the structure of **M**^-1^ in terms of the structure of **J** would allow us to connect the dynamics of the network to the kinetic properties of single reactions, providing insight into the origin of the network dynamic structure. Any numerical investigation of these properties should be performed on a real metabolic network, where network topology as well as order of magnitude differences in reaction fluxes, metabolite concentrations, and reaction rate constants are essential features in determining the dynamics of the network (22).

In this study, we present key results on the modal structure of kinetic models of metabolism, using the metabolic network in the human red blood cell (RBC) as an example (22). This model consists of ten enzyme mechanisms represented by mass action kinetics inserted in a background of 133 approximated rate law reactions (3)(23), parameterized with measured metabolite concentrations and enzyme kinetic constants. Using both numerical and theoretical arguments, we demonstrate how the dynamic structure of the modal matrix **M**^-1^ forms due to specific properties of the Jacobian (**J**) matrix. Using Gershgorin circle theorem, we first explain simple dynamic structures in cases where diagonal dominance occurs in **J**. Second, we use the matrix power iteration algorithm to show how modes with more complicated sparsity structures arise from topologically connected elements of **J** that have similar magnitude. Third, we describe how such complicated mode structures arise due to specific metabolite and reaction properties of the system, through examination of the corresponding elements in **S** and **G**.

We focus on demonstrating general principles through a set of case studies on the concentration Jacobian matrix and the mode structures associated with metabolite groups. These principles also apply to the flux Jacobian matrix and the relate flux modal structures, which are characterized in terms of the flux variables and describe the dynamic properties of the reaction groups (24).

## Materials and Methods

### Software

All work was done in Mathematica 10. We used a package called the MASS Toolbox (https://github.com/opencobra/MASS-Toolbox) for model simulation and analysis.

### Model simulation and perturbation

The model used in this study is a whole-cell kinetic model of red blood cell (RBC) metabolism with 10 enzyme modules incorporated (22). An enzyme module describes the detailed reaction steps of an enzyme catalyzed reaction, including substrate binding, catalytic conversion, product release and regulatory actions. The 10 enzyme modules are mainly located in glycolysis and the pentose phosphate pathway.

We used measured steady state metabolite concentrations as the starting state of the system before the perturbation. The perturbation used in this study was to simulate ATP hydrolysis in RBC. At time 0, the ATP concentration was decreased by 0.1 mmol/L while ADP and Pi concentrations were increased by 0.1 mmol/L. We then simulated the subsequent concentration and flux changes through numerical integration of the ODE equations. We gave the system enough time (10^6^ hours) to regain the steady state concentrations. The dynamic response of a specific metabolite or a combination of metabolites over time was visualized using the plotting functions in MASS Toolbox.

### Mode structure interpretation and dominant mode selection

To simplify the mode structure for interpretation, we neglected metabolites whose absolute coefficient values are less than 5% of the maximum absolute coefficient. We found that generally metabolites with small coefficients do not substantially contribute to the dynamic response of the mode, and 5% serves as a useful cutoff value for purposes of analysis.

When selecting modes that can be explained by diagonal dominance alone, we applied the following criteria to both concentration modes and flux modes. When examining a particular mode, we first neglected elements whose absolute coefficient values are less than 5% of the maximum absolute coefficient. If there is only one element left in the mode and it is diagonally dominant, the mode is explained by diagonal dominance. For modes with multiple elements, we selected the mode where its largest coefficient is at least twice as large as the next one and corresponds to the most diagonally dominant element in the mode.

### Power iteration and Hotelling’s deflation

Since the modes are left eigenvectors of the Jacobian matrix, we left multiplied the Jacobian matrix by the vector during power iteration. We started with a random vector, obtained a new vector after matrix multiplication and normalized against the Euclidean norm. We kept running this iteration until the length of the ending vector converges. The algorithm is demonstrated as follows,

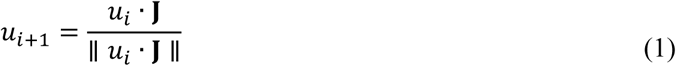

Where *i* is the number of iterations, *u*_*i*_ is the starting vector and *u*_*i*+1_ is the ending vector in each iteration.

Since power iteration only calculates the leading eigenvalue and eigenvector of the Jacobian matrix, we used Hotelling’s deflation to remove the impact of the leading eigenvector and calculated the next leading eigenvector (25). The algorithm thus results

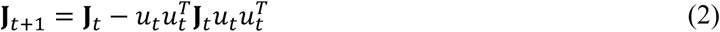

Where *J*_t+1_ is the Jacobian matrix after the leading eigenvector *u*_*t*_ of the previous Jacobian matrix *J*_*t*_ removed.

In cases where the eigenvalues are clustered together, different starting vectors will result in different eigenvectors at the end of iteration. To compare the approximated eigenvectors from power iteration with the actual eigenvectors, we picked the eigenvalue cluster with time scale around 0.016 milliseconds and reduced **J** using Hotelling’s deflation method until this time scale was reached. We started with 100 random vectors and multiplied them by **J** through 100 iterations, which we found to be large enough for the vector to converge in practical cases. To obtain the set of linearly independent vectors out of the 10^4^ vectors, we started with one of the vectors, added another vector (from the 10^4^ vectors), and calculated the rank of the matrix formed by the current vector space. We kept adding the vector one at a time for all the ones we calculated. If the matrix rank increases, the added vector is linearly independent with the earlier vectors and will be kept in the final vector set. Otherwise, it will not be included. We also calculated the norms of all vectors during iterations as eigenvalue approximations for comparison with the eigenvalue cluster.

## Results

### Linear analysis on dynamic structures of the metabolic network

We first briefly introduce the basic theory for linear analysis of metabolic networks. In a biochemical reaction network, the dynamic mass balances for all *m* concentrations **x** are given in the form of a matrix equation:

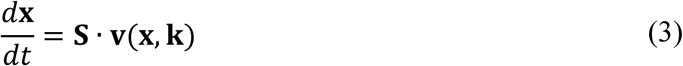

Where **S** is the *m* × *n* stoichiometric matrix, **x** is the *m* × 1 vector of metabolite concentrations, and **v** is the *n* × 1 vector of reaction fluxes. The formulation of **v** depends on the reaction rate law used and the mass action rate law is expressed as a function of the concentrations **x** and kinetic parameters **K**.

Linearizing around a particular steady state **x**_0_ (i.e., **S.v(x**_0,_, **k**)=0) yields,

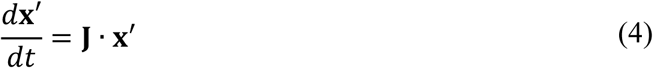

Where **x**^**′**^=**x−x**_**0**_are the concentration deviation variables from the steady state and **J** = **S · G** is the concentration Jacobian matrix (20). **G** (=*d***v**/*d***x**) is the gradient matrix obtained from linearization of the reaction rates (24). It is the same matrix as the non-normalized elasticity matrix from metabolic control analysis (19).

An eigen-decomposition of the Jacobian matrix yields a different representation of the same linearized system, with dynamically independent motions of metabolites grouped into modes within the modal matrix (20).

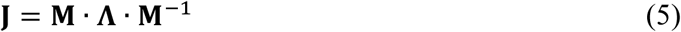

Where **M**^-1^ is the modal matrix and 
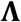
 is the diagonal matrix of eigenvalues. During eigen-decomposition, we can append the left null space vectors of the Jacobian matrix to the modal matrix and assign those vectors zero eigenvalues. This operation makes both modal matrices full rank since a rank deficient matrix is not invertible. The modes are defined as *m*= **M**^−1^ ∙**x**. Substituting Eq. 5 into Eq. 4, and based on the mode definitions, we have,

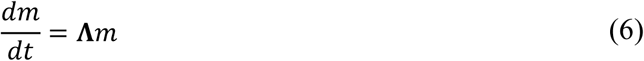

As defined in Eq. 6, the eigenvalues and modes give information on the dynamically independent motions of metabolite groups (20).

The rows of the modal matrix, which correspond to modes, are left eigenvectors of **J** (*u***J** = λ*u*). Each mode is associated with an eigenvalue and represents the dynamic motion in a characteristic time scale defined by the eigenvalue. These characteristic time scales describe the approximate time it takes for the mode to relax (return near its original reference state) when the system is perturbed from steady state (see Supporting Material). Our focus in this work is to examine the sparsity structure of the modes and determine how this structure is connected to properties of the Jacobian matrix.

### Half-reaction equilibria resulting from linearization of bilinear mass action rate laws are key dynamic features of **G**

Understanding the sparsity structure of the modal matrix **M**^**-1**^ derived from the Jacobian matrix **J** is greatly helped by the knowledge that **J** is sparse and contains elements spanning several orders of magnitude. The gradient matrix **G** (=*d***v/***d***x**) is responsible for the order of magnitude scaling of the elements of **J**, since **J** = **S · G**, and the elements of **S** are mostly -1, 0 or 1. For mass action reactions, the *d***v**/*d***x** derivatives comprising the gradient matrix **G** have a specific mathematical form and biochemical interpretation (Figure 1). The form of the mass action rate law for an example bilinear reaction between metabolite A and enzyme form E where 
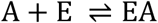
 is 
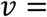
 and the three resulting *d*v/*d*x terms in **G** for the reaction are *k*^+^[A], *k*^+^[E], − *k*^−^. From these three terms, we can see that certain reactant/product terms are eliminated when calculating the reaction sensitivities (derivatives in the form of *d***v**/*d***x**) in **G**. This mathematical operation can be interpreted as splitting the original reaction into ***half reactions*** in a biochemical context. In the case of bilinear kinetics of enzymatic binding/release reactions, the half reaction describes the binding/release process for one reactant, which is held constant.

**Figure 1.**
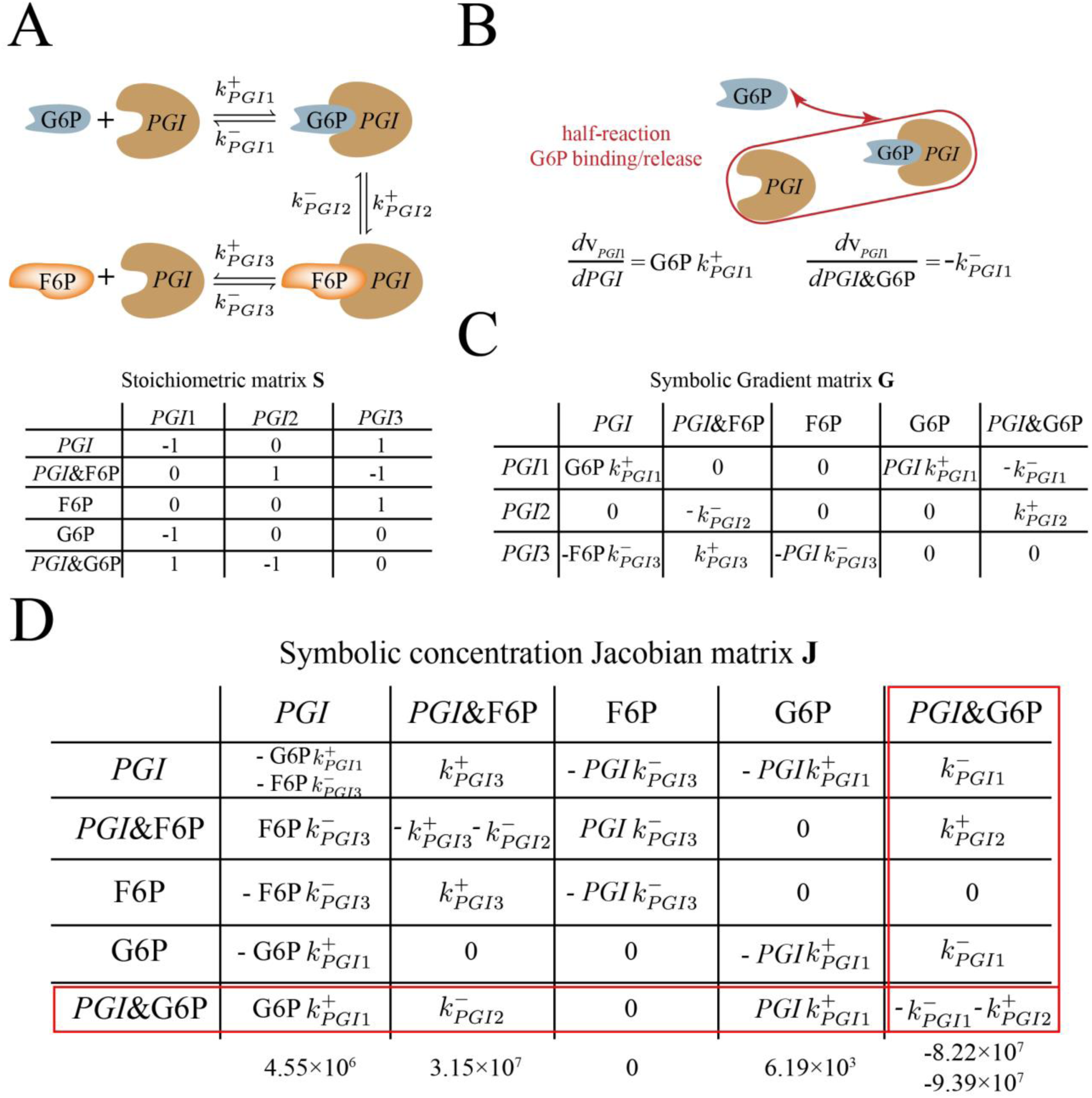
*PGI* enzyme module and its associated matrices. (A) A schematic diagram of individual reaction steps associated with *PGI* enzyme module and its stoichiometric matrix. The PGI enzyme module consists of three reaction steps: binding of G6P (*PGI*1), conversion of G6P to F6P (*PGI*2) and release of F6P (*PGI*3). The enzyme form *PGI* is in italic. We use an “&” notation to denote that the enzyme form is bound with metabolite(s). (B) Graphical representation of the concept of half reaction. Here we demonstrate the half reaction associated with the binding/release process of G6P, which is held constant. To determine the equilibrium state of this half reaction, we are comparing the sensitivities associated with *PGI* (G6PK^+^_*PGI*1_) and *PGI*&G6P (−K_PGI1_^−^). This comparison is equivalent to comparing G6P concentration and1/K_eq,PG1_. (C) The gradient matrix of the PGI enzyme module. The gradient matrix (=*d***v**/*d***x**) is obtained from linearization of the reaction rates and represents reaction sensitivities to metabolite concentrations. (D) The cause of diagonal dominance demonstrated through the symbolic concentration Jacobian matrix of the *PGI* enzyme module. Using row 5 as a case study, we observe that, in the case of mass action rate law, diagonal dominance is determined by the distance from half-reaction equilibrium for individual half-reactions. When comparing the terms associated with *PGI*1 reaction between diagonal and off-diagonal positions, we are comparing the sensitivity of G6P ( *PGIK_PGI1_^+^*) and sensitivity of *PGI* (G6PK_PGI1_^+^) with that of *PGI*&G6P −K_PGI1_^−^). This comparison is equivalent to comparing the concentrations of *PGI* and G6P with *K_d, 1_*(*K_d,–PGI1_* = *_KPGI1_^−^/ k_PGI1_^+^*), thus determining the distance from equilibrium for *PGI* and G6P binding/release half-reactions. The numerical values for each entry in row 5 is below the symbolic forms. Additionally, we can see clearly that column dominance cannot happen in the concentration Jacobian matrix due to the structure of mass action rate law. In the current case, we can see that the absolute sum of off-diagonal elements in a column is always at least as large as the absolute diagonal element, meaning that diagonal dominance does not occur across columns.

In a biochemical network, the concentrations can span different orders of magnitude, resulting in their activities at different time scales in response to perturbations. Typically, the enzyme forms have much smaller concentrations than those of metabolites. For these reactions, the half reactions will have sensitivities on different orders of magnitude, separating the dynamics associated with the enzyme forms apart from those of metabolites. In case of the bilinear reaction mentioned above, we observe such half reaction is associated with the binding/release of A, which is relatively constant due to its larger concentration. On the other hand, E and EA form a quasi equilibrium due to their smaller concentrations.

For a full reaction, the distance from equilibrium is defined as Γ/K_eq_, where Γ is the mass action ratio and K_eq_ is the equilibrium constant. Thus, for the example bilinear reaction mentioned above, its distance from equilibrium can be expressed as *k*^−^[EA]/ *k*^+^[A][E]. Similarly, the distance from equilibrium for the half reaction associated with binding/release of A can be expressed as the ratio between the reaction sensitivities of E (*k*^+^[A]) and EA (*k* −). This ratio can be simplified into [A]/Kd, A, where Kd, A equals *k*−/ *k*^+^ and represents the dissociation constant for binding/release of A. In cases where there is only one reactant on both sides of the reaction, the half-reaction equilibrium is equivalent to the equilibrium of the reaction itself (since the resulting dynamic ratio is *k*^+^/ *k*^−^).

As a specific example, we present a case study on the glucose 6-phosphate isomerase (*PGI*) enzyme module (Figure 1A) from a whole-cell kinetic model of RBC metabolism (22). An enzyme module describes the individual reaction steps of an enzyme-catalyzed biochemical reaction, and each step is represented by a mass action rate law. Using *PGI*1 reaction as an example, the half reaction of interest is the binding/release of glucose 6-phosphate (G6P) (Figure1B red). The comparison of the sensitivities of *PGI* (G6P _k_+_PGI1_) with *PGI*&G6P (− *k*^-^_PGI1_) (& denotes *PGI* bound with metabolite G6P) in magnitude is equivalent to the comparison of G6P concentration with 1/K_eq,_ PGI1_1_. This comparison effectively results in determining the distance from equilibrium for G6P binding/release half reaction. It is worth noting that the full equilibrium ratio would include the enzyme forms that have been removed by differentiation and therefore do not influence the above comparison; thus, the distinct definition of a half-reaction equilibrium ratio is helpful.

Understanding how mode structure forms requires an understanding of the relationships between terms in **J**, which in turn are set by particular reaction sensitivities from **G**. As we will show later, the complexity of a mode is dependent on the distance from equilibrium of half reactions defined by these sensitivities in **G**. Half reactions that are far from equilibrium result in simple mode structures while those near equilibrium together form complex modes.

### Diagonal dominance and the Gershgorin circle theorem applied to the Jacobian matrix

Now that the biochemical origin of terms in the key matrices **G** and **J** are clarified, we move to a discussion of modal structure in kinetic models of metabolism. The modes are defined by

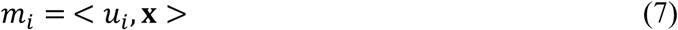

Where *u_*i*_* the left eigenvector and **x** is the steady state concentration vector. The relative magnitudes of the elements of *u_*i*_* determine the effective sparsity of a mode when low contributing elements are truncated. However, since the modes are calculated through a numerical algorithm, it is usually not straightforward to link a mode composition to particular elements of the Jacobian matrix, unless the Jacobian matrix has certain structural properties. One such property is diagonal dominance of the rows or columns of the Jacobian, which occurs when the magnitude of a diagonal element is greater than the sum of the magnitudes of off-diagonal elements in the same row (in the case of row dominance) or column (column dominance), see Figure 2A. We focus on row dominance in this work, as column dominance does not occur in the concentration Jacobian matrix due to the structure of the mass action rate law, as demonstrated in Figure 1D.

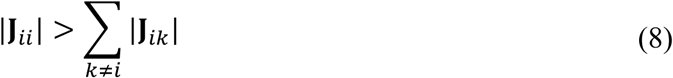

The degree of diagonal dominance of a row number *i* can be quantitatively described by a metric we term the diagonal fraction, defined as the ratio between the sum of the absolute values of off-diagonal elements and the absolute value of the diagonal element:

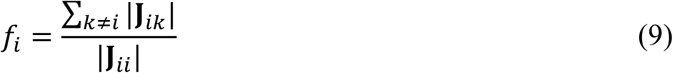

**Figure 2.**
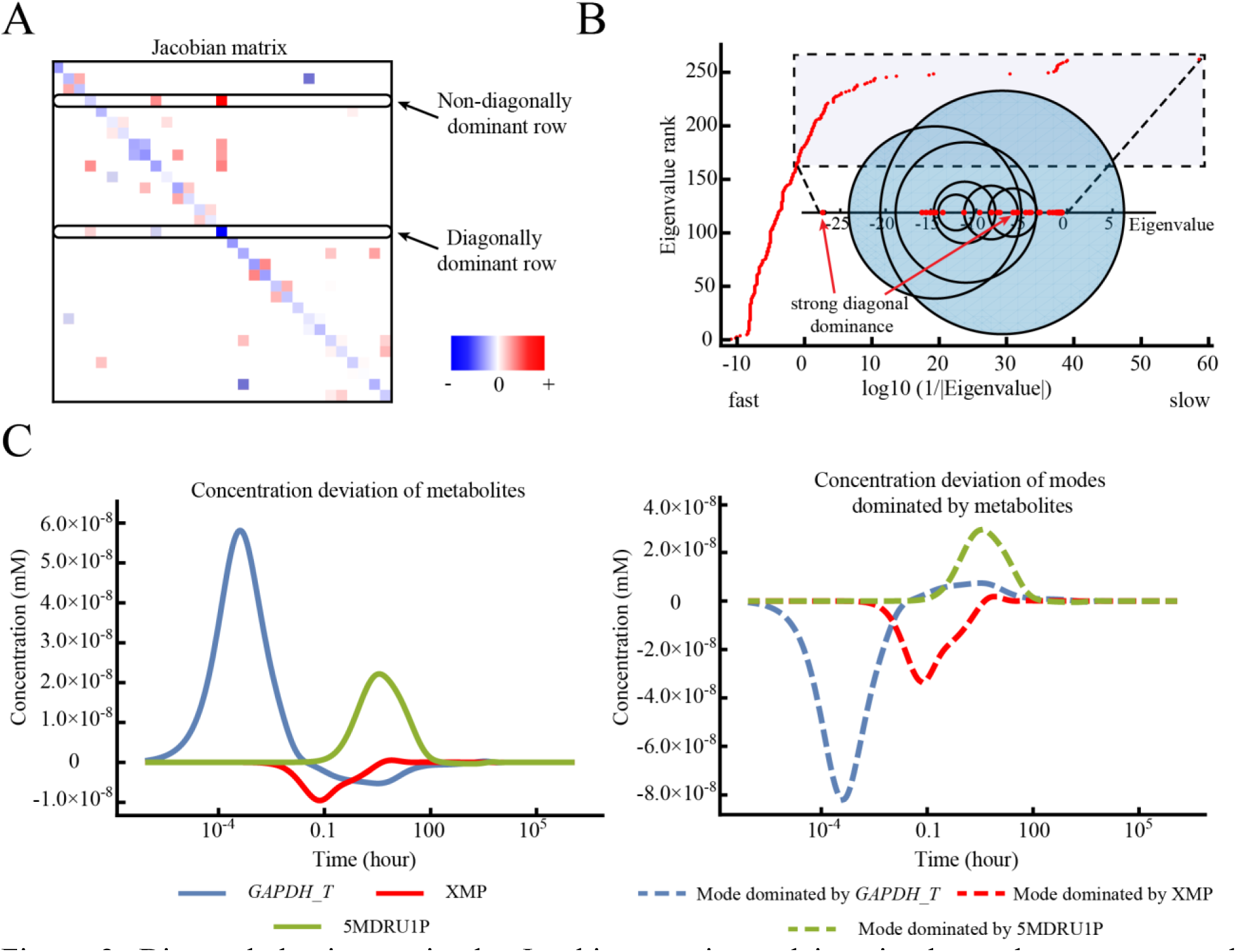
Diagonal dominance in the Jacobian matrix explains simple mode structures and corresponding eigenvalues with the help of Gershgorin circle theorem. (A) Example Jacobian matrix of the RBC metabolic network (22) with different degrees of diagonal dominance. The Jacobian matrix of the metabolic network has a sparse structure, and the diagonal elements of the matrix are always negative due to the structure of the rate laws used. The matrix was extracted from the full concentration Jacobian matrix for illustrative purposes. (B) The entire set of eigenvalues of the Jacobian matrix is shown in the larger plot, with x-axis denoting the inverse of absolute eigenvalues at the log10 scale. In the inset, selected Gershgorin circles of the Jacobian matrix with circle centers ranging from -27 to -5 are shown for illustrative purposes. Eigenvalues greater than -27 are drawn together with the selected circles. The Gershgorin circles from rows with strong diagonal dominance have centers at -26.2 and -5.26 as shown, and the eigenvalues inside are -26.3 and -5.33. All eigenvalues are negative as the system is dynamically stable. The imaginary components of the eigenvalues are small and therefore are neglected. (C) The dynamic response of *GAPDH_T*, XMP, 5MDRU1P, compared to the respective modes dominated by these metabolites/enzymes, under an ATP hydrolysis perturbation. The dynamics of the mode dominated by a single metabolite coincide with the dynamics of that metabolite. These modes occur at fast, intermediate and slow timescales, showing that diagonal dominance can occur at any time as long as the structural properties of the Jacobian matrix allow.

Diagonal dominance of a row of the Jacobian matrix gives information about its corresponding eigenvalue. This relationship is made clear using Gershgorin’s circle theorem (26), which constrains an eigenvalue to be within a certain radius, based on the sum of the off-diagonal elements in a particular row/column, of the diagonal element. The theorem is particularly useful in confining eigenvalues within Gershgorin circles when strong diagonal dominance (a small *f*_*i*_ value) occurs, as the eigenvalue will be close to the diagonal element of the dominant row.

### Diagonal dominance in the Jacobian matrix underlies simple mode structures

To demonstrate the occurrence and impact of diagonal dominance in a real metabolic network, we use the RBC kinetic model to draw the Gershgorin circles and the eigenvalues from **J** (Figure 2B along x-axis). As highlighted in Figure 2B, for the selected set of Gershgorin circles, there are two cases where the circle resulting from the strongly diagonally dominant row is very constrained and a unique eigenvalue falls inside the circle. In those cases, the eigenvalue is very closely approximated by the diagonal element.

In addition to providing information about the eigenvalues, diagonal dominance in **J** also causes a simple sparsity structure within modes corresponding to these eigenvalues. When a row has strong diagonal dominance (*f* < 0.1), the diagonal metabolite usually is the only significant nonzero element in the mode (Table S1). For example, the enzyme form *GAPDH_T* (glyceraldehyde 3-phosphate dehydrogenase at tense state) has a very small diagonal fraction value, and is the only element in the mode at its corresponding time scale. The underlying reaction that causes its dominance is the transition step from enzyme form *GAPDH* at relaxed state to tense state *GAPDH* ⇌ *GAPDH_T*, where the sensitivity of *GAPDH_T* (−***K***_GAPDH___transition_step_) contributes the most to its diagonal element in **J**. When a mode contains only the diagonally dominant metabolite, the dynamic motion of the mode drives that metabolite back to its reference state on a timescale determined by the eigenvalue. For example, under ATP hydrolysis perturbation, the dynamics of GAPDH_T match closely with the dynamics of the mode in which *GAPDH_T* is dominant (Figure 2C). When diagonal dominance becomes weaker (*f* > 0.1), the diagonally dominant metabolite shares modes with other metabolites, as demonstrated in the case of enzyme form glucose 6-phosphate dehydrogenase bound with 6-phospho-D-glucono-1,5-lactone (*G6PDH*&6PGL) in Table S1. In those cases, the ratio between those metabolites in the mode is similar to that in the diagonally dominant row of the Jacobian matrix. Overall, in the RBC metabolic model used in this work, the structure of 38 out of 244 (15.6%) concentration modes can be explained by diagonally dominant metabolites. Other statistics about diagonal dominance in rows of concentration Jacobian matrix can be found in Table S2 and Figure S1.

As another effect of diagonal dominance, there exists an important relationship between diagonal dominance in **J** and system dynamic stability, which is characterized by the sign of eigenvalues of **J** in that any positive eigenvalues result in the steady state being unstable. Negative diagonal elements in **J** strongly support system stability, and this effect is further magnified by diagonal dominance (see Supporting Material and Figure S2).

### Dependence of diagonal dominance on the parameters of the metabolic network

Having established the usefulness of diagonal dominance for understanding eigenvalue and mode structure, we now describe the origin of diagonal dominance in terms of the parameters of the system. To understand how diagonal dominance in **J** is manifested through reaction properties, we can examine the association of elements between **J** and **G**. We can see that for each diagonally dominant metabolite (diagonal fraction < 1), its diagonal element in **J** can be matched with a specific reaction sensitivity element for that metabolite similar in absolute value in **G**. Such an element is the largest in absolute value for the flux-concentration derivatives (d**v**/d**x**) associated with that metabolite. Therefore, a single term in **G** dominates the resulting diagonal term in **J** (Figure S3). Furthermore, single reaction sensitivities in the form of d**v**/d**x** in **G** can determine the dynamic behavior of the system in terms of the resulting eigenvalues when diagonal dominance occurs. This correspondence can also be extended to metabolites with non-diagonal dominance (Figure S3), indicating the interpretable connection between **J** and **G**.

As a case study, we examine the cause of diagonal dominance in **J** of the *PGI* enzyme module. We see that, in the enzyme module, diagonal dominance in **J** is determined by a particular half-reaction equilibrium ratio, as defined above. We demonstrate this by examining the enzyme form *PGI*&G6P in the 5^th^ row of **J** (Figure 1D). The diagonal term of **J** for *PGI*&G6P shows that the enzyme form is associated with two reactions, *PGI*1 and *PGI*2. Specifically, reaction *PGI*1 can be split into two half reactions, related to G6P binding/release and PGI binding/release processes. The comparison of the diagonal term (−*K^−^_PGI1_*) with the off-diagonal terms (G6PK^+^_*PGI1*_ and *PGIK*^+^_*PGI1*_) related to *PGI*1 reaction is effectively examining the associated half-reaction equilibrium ratios, which are G6P/*K_d,PGI1_* and PGI/*K*_d,*PGI*1_(K_d,*PGI1*_=K^−^_*PGI 1*_/*K*^+^_*PGI1*_)The term G6PK_PGI1_^+^ is smaller than −*k*_*PGI*1_^−^ on the diagonal position in magnitude while _*PGIK*_^+^*PGI*1 term is negligible compared to − *k*_*PGI*1_^−^, due to the small concentration of the *PGI* enzyme form. For reaction *PGI*2, the term *K*_*PGI*2_^+^ at the diagonal position is much greater than *K*^-^_*PGI*2_, with the consumption of *PGI*&G6P favored. As a result, the diagonal term of **J** for *PGI*&G6P is greater than the sum of off-diagonal terms in the same row, resulting in diagonal dominance.

To summarize, diagonal dominance can be understood based on the distance from half-reaction equilibrium, by comparing metabolite concentrations to the reaction equilibrium constant. In the case of a single reactant on each side of the reaction, the equilibrium constant alone affects the degree of diagonal dominance. This type of analysis can also be applied to other enzyme forms in **J**.

### Power iteration connects mode structure to the structure of the Jacobian matrix

Diagonal dominance explains the structure of most of the highly sparse modes, but cannot address mode structures that are complicated by more than one or two significant elements. We now show how more complicated mode structures form mathematically from specific elements of the Jacobian matrix. We demonstrate that examining the modes of the Jacobian matrix from the perspective of the matrix power iteration algorithm is illustrative in describing how complicated mode structures arise.

Matrix power iteration is an algorithm to calculate the leading eigenvalue and eigenvector of a matrix (or left eigenvectors in the case of the modes) (27). In the power iteration algorithm, the Jacobian matrix is left multiplied by a random vector (*u*i), the resulting vector is normalized, and this process is repeated until the vector converges. (Figure 3A). If the eigenvalue with the largest magnitude is well separated from the other eigenvalues;, the final vector will converge to the corresponding leading eigenvector. The Euclidean norm of *u***J** in the last iteration will be the associated leading eigenvalue λ, where *u***J** =λ *u*. During the iteration process, the elements of the Jacobian matrix that contribute to the modes will “stretch” the vector through multiplication in the direction of the leading eigenvector. Given the fact that the Jacobian matrix is sparse, the power iteration algorithm can help us understand eigenvector structure by inspecting how the Jacobian elements stretch the vector to ultimately result in the eigenvector.

**Figure 3.**
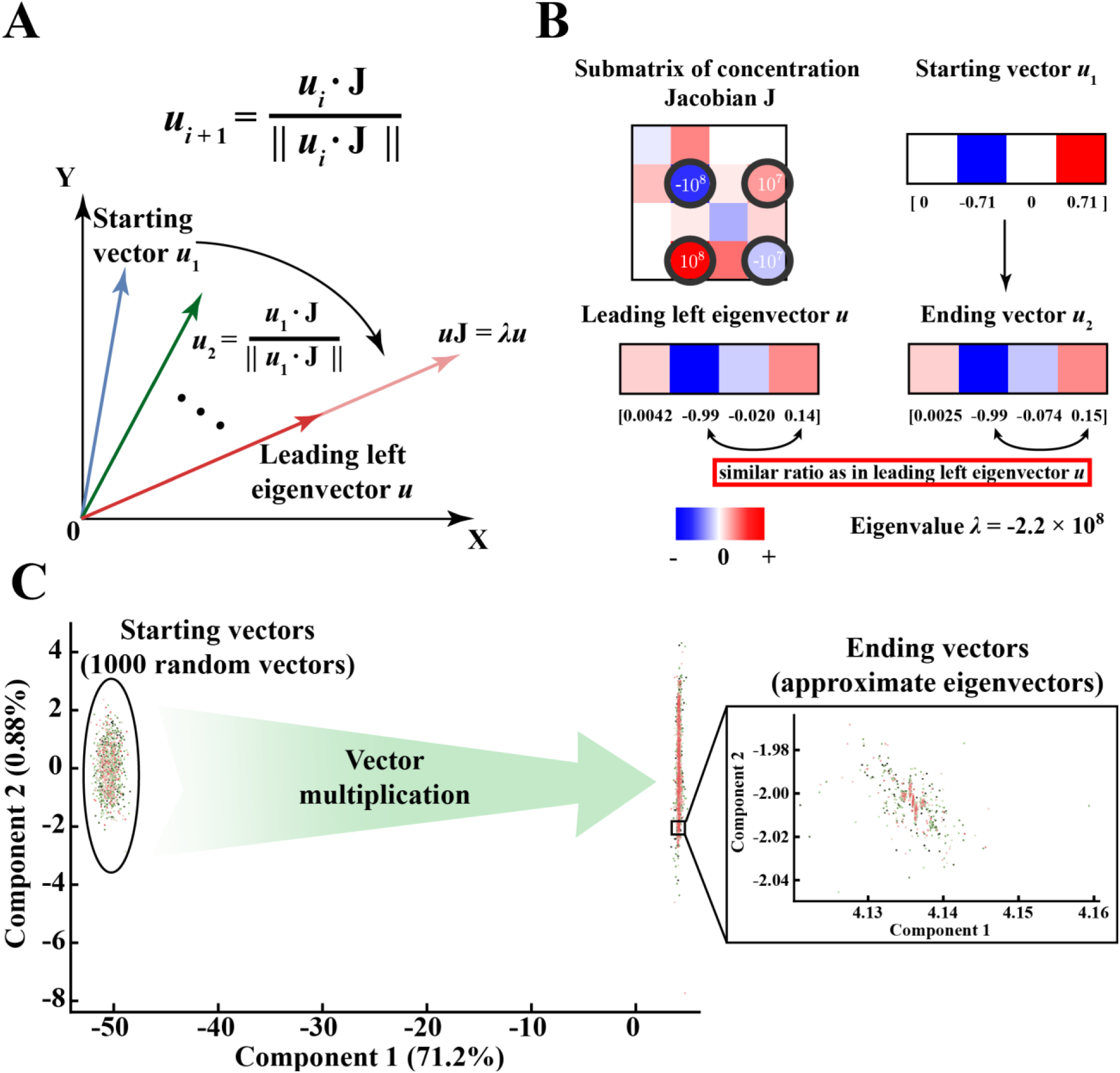
The power iteration algorithm demonstrates how complicated dynamic structures arise from topologically connected elements of similar magnitude within the Jacobian matrix. (A) Power iteration can be used to calculate the dominant left eigenvector of the Jacobian matrix. The left eigenvectors are the modes of the metabolic network. The algorithm left multiplies the Jacobian matrix by a random vector (*ui*), normalizes the resulting vector and repeats the process until the vector converges to the eigenvector. (B) Topologically connected Jacobian elements of similar magnitude determine complicated eigenvector structure. In this case study, we extracted a submatrix of **J** that corresponds to the nonzero elements of a certain eigenvector, which contains *G6PDH* enzyme forms. The four Jacobian elements (also the largest) that are key in determining this eigenvector structure are located in the 2nd and 4th rows, circled in black. Specifically, the structure of 2nd or 4th rows matches closely with that of the eigenvector, with similar ratios at the 2nd and 4th positions. Multiplying the Jacobian matrix by any non-orthogonal starting vector (*u*1), for example the one shown, results in a vector (*u*2) that has a structure more similar to the eigenvector. The contribution of those rows individually to eigenvector formation are further shown in Figure 4 and S4. For clear demonstration purposes, the comparison of relative colors only works for individual box (surrounded by black stroke) itself, but not across different boxes. (C) Principal component analysis on all power iteration vectors starting with 1000 different random vectors. We randomly picked 1000 starting vectors and multiplied them with the full Jacobian matrix (292 × 292). The starting vector is multiplied through several iterations (10 ~ 20) until it converges to the eigenvector (the dot product of the ending vector and the eigenvector is no greater than 1.0001 and no less than 0.9999). We then performed principal component analysis on all iteration vectors (including the starting vectors) and plotted each vector in terms of the contribution from the first two principal components. The first principal component corresponds to the leading eigenvector of the Jacobian matrix while the rest of components (less than 1% contribution each, only component 2 shown here) together explain the variation of the vector from the eigenvector. Ideally, the contribution of the rest of components will be 0 when the ending vector becomes the eigenvector. However, due to large order of magnitude differences between elements in **J** and the cutoff we set when comparing the ending vector with the eigenvector, we ended up with variations from the eigenvector (nonzero contribution of component 2 in the inset plot).

To illustrate the process of vectors converging to the leading eigenvector through power iteration, we perform power iteration algorithms on 1000 random starting vectors using the full Jacobian matrix (292 × 292). We then perform principal component analysis (PCA) on all the iteration vectors (Figure 3C). The random starting vectors quickly converge in the dimension of the first principal component (71.2% contribution), representing the eigenvector, and stabilize in the dimension of the rest of components (second principal component shown only, contributing a very minor percentage) after around 10 to 20 iterations.

As a technical detail of the implementation, a limitation of the power iteration algorithm is that it only calculates the leading eigenvalue and eigenvector. To calculate the next largest eigenvalue and the associated eigenvector, we must modify **J** to eliminate the impact of the previous eigenvector and eigenvalue at each step. Such elimination can be accomplished with the Hotelling deflation method (25), which returns a modified **J**, with the leading eigenvector and eigenvalue removed, that can be used for a new round of eigenvector and eigenvalue calculations using power iteration (see Materials and Methods).

### A case study on using power iteration to understand complicated mode structure

We use the power iteration method to demonstrate how the eigenvectors with more complicated structures form. In this section, we show that that the topological connection of elements of similar orders of magnitude in **J** is critical in determining the sparsity structure of the eigenvectors. This similar order of magnitude tends to lie around the eigenvalue (Figure 3B).

As a case study, we extract a submatrix of **J** (4 × 4) corresponding to the positions of nonzero elements (see Materials and Methods for cutoff) of a particular eigenvector, which is associated with G6PDH enzyme forms of the RBC metabolic network. When **J** is pre-multiplied by a pseudo-random starting row vector, we see that the ending vector matches closely with the actual eigenvector (Figure 3B and S4A). It is clear upon inspection that the largest values in the submatrix are also the largest values in the mode. The four key **J** elements (also largest in the submatrix) determining eigenvector formation are located in the 2^nd^ and 4^th^ rows (Figure 3B black circles). These rows both have similar structures to the eigenvector, where the ratio between the 2^nd^ and 4^th^ elements in the row is the same as that in the eigenvector. This shows that the matrix structure is reflected in the eigenvector structure.

To explore how the 2^nd^ and 4^th^ rows both contribute to eigenvector formation, we can perturb the starting vector such that it interacts with these rows specifically, such as (0, -1, 0, 0) and (0, 0, 0, 1), to examine each row’s effect individually. As a result, starting from either vector leads to a structure similar to the original eigenvector (Figure S4B and S4C). Thus, it seems that both rows have similar contributions to the structure of the eigenvector in this case, although their magnitude is different. Together, the four elements in those two rows (Figure 3B black circles) form a topologically connected structure and interact with each other symmetrically to determine the eigenvector structure. The other large element at position (4, 3) is not involved with this symmetric interaction and thus has a smaller contribution to eigenvector formation.

Next, to demonstrate the interplay of the submatrix elements, we show how modifying the four key elements of the sub-matrix changes the eigenvector. First, to examine the impact of the largest diagonal element in the submatrix at position (2, 2), we modify the diagonal element at position (4, 4) to have the same value as the element at (2, 2) (Figure 4B). The resulting vector has a different ratio between its elements compared to the original **J** eigenvector, with a larger value in the 4^th^ element, reflecting the larger value in the (4, 4) position of the submatrix. We then further change the off-diagonal element of **J** at (2, 4) to be the same as the element at (4, 2) to create a more symmetric structure (Figure 4C). The resulting vector now has the same value on both the 2^nd^ and 4^th^ positions, showing that the off-diagonal elements modify the weightings on the eigenvector, and a fully symmetric Jacobian structure will result in an equally weighted eigenvector structure. These perturbations show that how the relative values of the dominant elements in a submatrix are clearly reflected in the corresponding mode structure.

**Figure 4.**
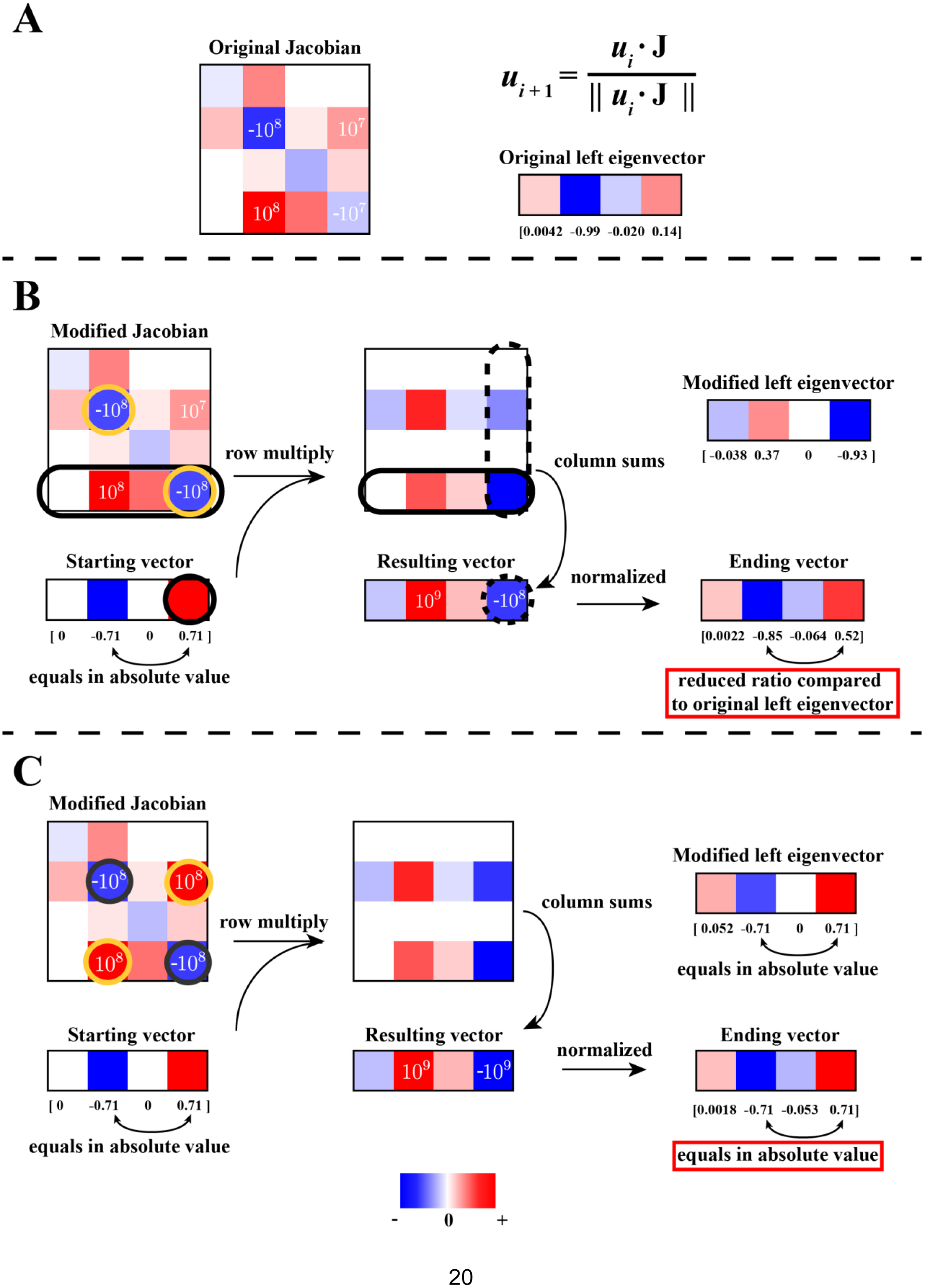
Figure 4. Analysis of complicated mode structure through power iteration with modified Jacobian matrix. We divide the vector multiplication with the Jacobian matrix into multiple steps. First of all, each row of the Jacobian matrix is multiplied by every element of the starting vector (Panel A solid black circles). We then sum up each column of the second matrix to obtain the resulting vector (Panel A dash black circles), which is normalized to give the ending vector. (A) The original Jacobian matrix and its leading left eigenvector. The matrix and the eigenvector are the same as in Figure 3 and will be used for comparison with later panels. (B) Starting vector multiplied with the modified Jacobian matrix. We modified the Jacobian element at position (4, 4) to be the same value as the element at position (2, 2). The ending vector has a smaller ratio between the 2nd and 4th elements than that of the original eigenvector, as would be expected with a larger absolute value at position (4, 4). The eigenvector of this modified matrix is shown in the upper right of the panel. (C) Starting vector multiplied with a different modified Jacobian matrix. We further changed the modified Jacobian matrix in panel A to create a more symmetric structure, where the element at position (2, 4) is same as the element at position (4, 2). The ending vector has the same absolute values at the 2nd and 4th positions, showing that a fully symmetric Jacobian structure will create an equally weighted structure in eigenvector. The eigenvector of this modified matrix is shown in upper right. Overall, we demonstrate that changing the Jacobian element at either diagonal or off-diagonal position can alter the eigenvector of the matrix in a predictable manner, based on the topological pattern of the key elements determining the eigenvector structure. For clear demonstration purposes, the comparison of relative colors only works for individual box (surrounded by black stroke) itself, but not across different boxes.

The power iteration algorithm is a useful tool to analytically understand the structure of complicated eigenvectors of a real system. We have demonstrated that the modes form from a network of topologically connected values of similar magnitude in the Jacobian matrix, and the relative ratio between these values influences the structure of the eigenvector. These trends, where an eigenvector can be linked to particular topologically-connected elements of **J** of similar magnitude, are generally applicable beyond this case study (Figure S5). The Jacobian modifications demonstrate that the eigenvector of the matrix can be altered in a predictable manner by changing either diagonal or off-diagonal Jacobian elements along the same order of magnitude.

### Complicated mode structure arises from connected reactions with similar dynamic sensitivities in **G**

Power iteration helps to show numerically how complicated modes arise due to particular structures in **J**. For metabolic networks constructed with mass action rate laws, these numerical values have clear biological interpretations. Next, we describe the origin of complicated mode structure in terms of specific metabolite and reaction properties of the system. Given the fact that **J** can be decomposed into **S** and **G**, which represent network topology and reaction sensitivities, respectively, we focus on the specific elements in **S** and **G** that give rise to the structure of **J** and the resulting mode structure.

We use the same case study presented in the previous section, regarding the mode and submatrix of **J** for *G6PDH* enzyme forms. The mode contains four *G6PDH* enzyme forms (red circles in Figure 5A), with *G6PDH*&6PGL and *G6PDH*&NADPH&6PGL being the most dominant elements. The mode structure is largely determined by the sensitivities of reaction 6 in **G** (*K*_6_^+^, NADPH *K*_6_−) (Figure 5C). This reaction releases NAPDH and its elements in **G** dominate the topologically connected **J** elements at positions (2,2), (2,4), (4,2) and (4,4) (Figure 5D). The two most dominant mode elements mentioned above are associated with reaction 6. Their corresponding **J** elements contain *K*_6_^+^ and NADPH *K*_6_^−^, which are close numerically, meaning that NADPH concentration is similar to the equilibrium constant of the half reaction for NAPDH binding/release, where the term ‘half reaction’ is used as defined above. The ratio between NADPH *k*_6_^−^ and *k*_6_^+^ (NADPH/*K*_d,6_, where K_d,6_ = K _eq,6_) defines a half-reaction equilibrium ratio that is the key in determining the eigenvector structure. If NADPH concentration is higher, reaction 6 will become more sensitive to the concentration of the released form *G6PDH*&6PGL, compared to that of bound form *G6PDH*&NADPH&6PGL. This change will cause enzyme form *G6PDH*&6PGL to become more dominant in the mode, due to its greater diagonal dominance in **J**. Additionally, reaction 7 has the same order of magnitude sensitivity in the forward direction (*K*_7_^+^) as reaction 6, but has a much smaller sensitivity when interacting with *G6PDH*&NADP&G6P in the reverse direction, thus resulting in a much smaller contribution to this enzyme form in the mode. Finally, the unbound *G6PDH* enzyme form, although topologically connected to other enzyme forms through reaction 4, is not prominently featured in the mode, since its sensitivities in **G** are at a smaller order of magnitude.

**Figure 5.**
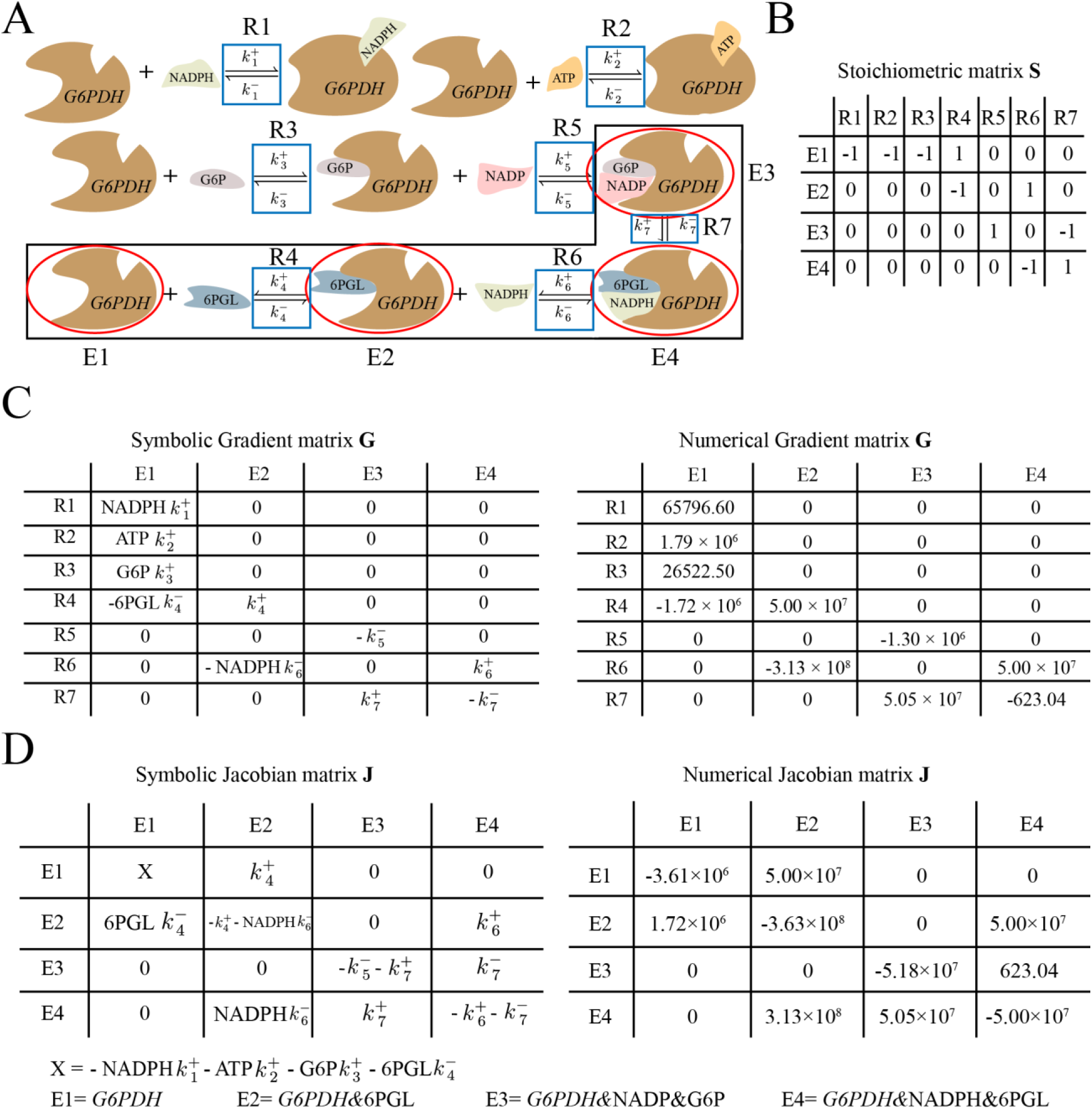
The origin of complicated mode structure associated with *G6PDH* enzyme forms demonstrated through the associated matrices. The mode structure contains four enzyme forms (denoted as E1, E2, E3 and E4, full annotation at the bottom), with *G6PDH*&NADPH&6PGL and *G6PDH*&6PGL being the most dominant elements. We extracted the submatrices associated with those four enzyme forms and their related reactions. We show that three key reactions and their associated reaction sensitivities in **G** determine the mode structure. (A) The reaction steps for the biochemical reaction catalyzed by *G6PDH* enzyme. The four dominant enzyme forms in the mode are labeled with red circles. The reactions with their notations (R1 to R7) are labeled with blue rectangular boxes. The three key reactions determining the mode structure are circle with black rectangular boxes. (B) The stoichiometric matrix **S** for the four enzyme forms in the mode and their associated reactions. The **S** matrix describes the network topology of the enzyme forms and determines how they interact in the Jacobian matrix. (C) The symbolic and numerical gradient matrix **G** for the four enzyme forms in the mode and their associated reactions. The key reaction sensitivities determining the two largest elements in the mode are associated with reaction 6 and its corresponding enzyme forms. The key terms are *k*^+^_6_ and NADPH *k*^−^_6_, which are similar in magnitude, due to the fact that NADPH concentration is similar to the equilibrium constant of the half reaction for NAPDH binding/release. (D) The symbolic and numerical Jacobian matrix **J** for the four enzyme forms in the mode. We found that the elements of reaction 6 in **G** dominate the topologically connected Jacobian elements that determine the mode structure. These elements are located at positions (2,2), (2,4), (4,2) and (4,4). Reaction 6 is connected to reaction 4 and 7, whose reaction sensitivities are much smaller in magnitude compared to that of reaction 6, resulting in very small coefficient for their associated elements in the mode (*G6PDH* and *G6PDH*&NADP&G6P).

Overall, only a few reaction sensitivities in **G** contribute to the mode structure in this case study, thus allowing us to determine the specific reactions that control the dynamics of the mode. For significant elements in the complicated mode structure, the associated half-reaction equilibrium constant is close to the metabolite concentration, thus creating dynamic interplay between multiple elements in the reactions. On the other hand, in the case of simple mode structure governed by diagonal dominance, the half-reaction equilibrium ratio associated with the diagonal metabolite is usually far from equilibrium. The analysis approach presented exploits the fact that dynamic features in **J** are an integration of the features in **S** and **G**, thus allowing us to understand modal structure in terms of both reaction sensitivities in **G** and network topology in **S**.

### Power iteration converges to eigenvector subspaces when eigenvalues are similar in magnitude

As an important technical aside, we note that the power iteration procedure works well when the eigenvalue is much larger in magnitude than the others; however, special behaviors arise when eigenvalues do not separate well. Specifically, when we reach modes where eigenvalues are close in magnitude, the power iteration algorithm converges to different ending vectors depending on the starting vectors. In this case, the starting vector is influenced by multiple eigenvectors comprising a subspace of dynamics active around this time scale, making the ending vector difficult to predict. The ending vectors overlap significantly with an “eigenvector subspace” (Figure 6A), as these vectors are influenced by multiple eigenvectors simultaneously. Also, the approximated eigenvalues overlap significantly with the actual eigenvalue cluster (Figure 6B), showing that the approximated eigenvalues settle in the range of the set of similarly leading eigenvalues. Overall, this analysis demonstrates how multiple eigenvectors influence dynamic response for time scales that are associated with multiple eigenvalues at similar magnitude.

**Figure 6.**
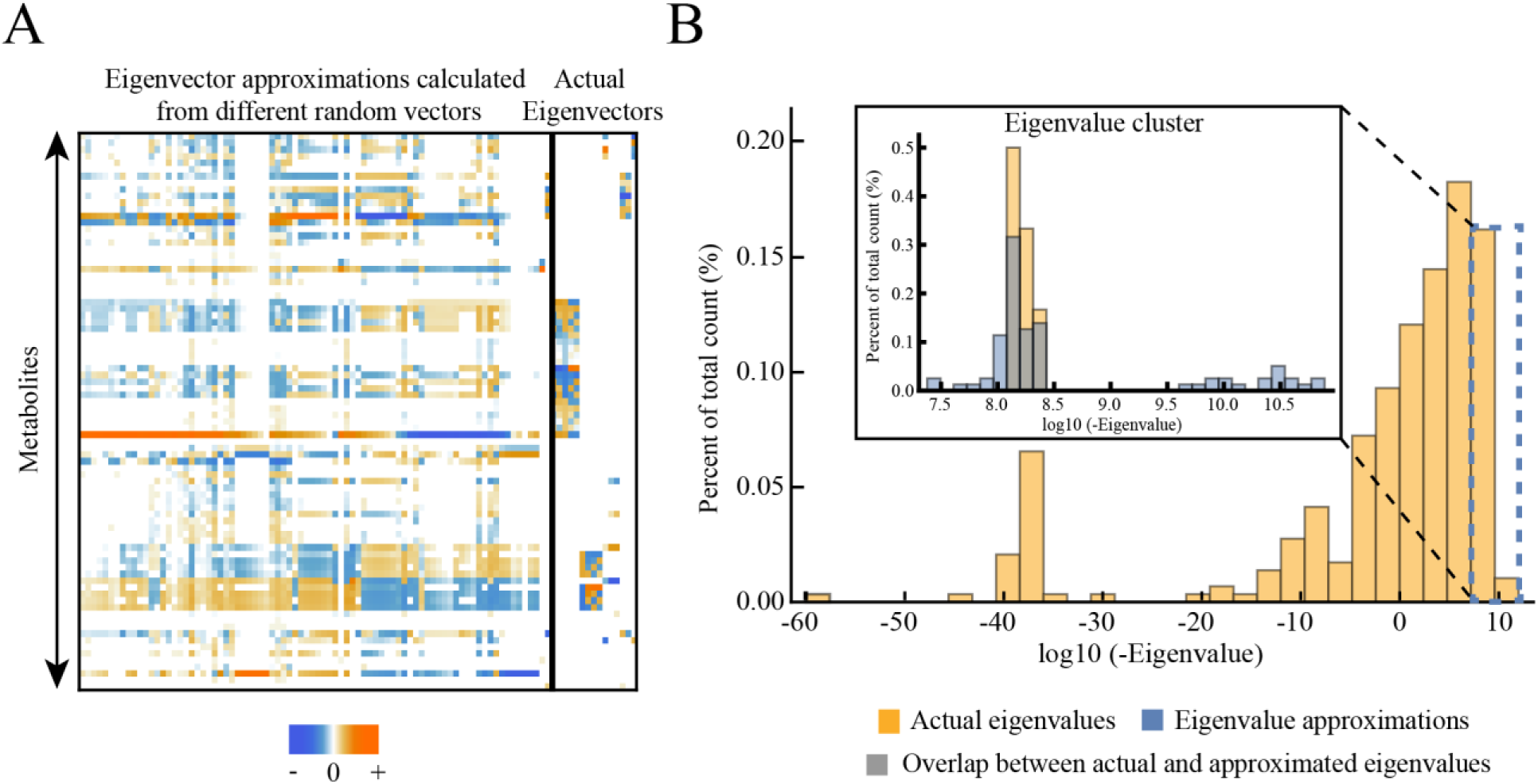
Eigenvalue and eigenvector approximations calculated from power iteration in cases where eigenvalues do not separate well. We selected a cluster of close eigenvalues (with a time scale around 0.016 milliseconds), reduced **J** using Hotelling’s deflation method until this time scale was reached (see Materials and Methods), and calculated approximated eigenvalues and eigenvectors using power iteration with different starting vectors. (A) Eigenvector approximations calculated during power iteration from different starting vectors, compared to the actual eigenvectors with eigenvalues in the selected range. We calculated the approximated 100 eigenvectors from 100 different random vectors with 100 iterations each and obtained vectors that are linearly independent with each other (see Materials and Methods). The left part of the matrix shown is the eigenvector approximations while the right part of the matrix shown is the actual eigenvectors, separately by the black bold vertical line. We found that the subspace formed by eigenvector approximations overlaps significantly with the actual eigenvector subspace. (B) The selected eigenvalue cluster is compared to the eigenvalue approximations calculated from power iteration. The selected eigenvalues and eigenvalue approximations are shown in the inset plot. We obtained the eigenvalue approximations from the same set of power iterations performed in panel A. The cluster of eigenvalue approximations overlaps significantly with the cluster of actual eigenvalues, showing that the eigenvalue approximations settle in the range of the set of similarly dominant eigenvalues.

## Discussion

In this study, we developed an understanding of how the dynamic properties of kinetic models of metabolism are reflected in their mode structures and how these structures are linked to specific properties of mass action reaction rate laws. 1) We showed that the diagonal dominance in rows of the Jacobian matrix is a frequent occurrence, and this feature results in simple mode structures where single metabolites relax back to their references states driven by particular eigenvalues. 2) For more complicated mode structures, we used the power iteration algorithm to show that these mode structures form from topologically connected values of similar orders of magnitude in the Jacobian matrix. 3) We showed that a key feature underlying mode structure is the reaction sensitivities in the gradient matrix **G**, which can be interpreted as the distance from equilibrium of half reactions defined by linearization of bilinear mass action equations.

Diagonal dominance of the Jacobian matrix as described by Gershgorin circle theorem gives information about certain eigenvalues. This property results in simple mode structures, which can occur on time scales that span different orders of magnitude. A simple structure dominated by a single element indicates that the concentration variable relaxes to its reference state after its characteristic timescale and does not interact with others on this timescale. Gershgorin circle theorem also has previously been applied to the Jacobian matrix of metabolic networks with a focus on examining system stability (28)(29).

We have shown that topologically connected elements of the Jacobian matrix at similar magnitude underlie complex mode structures. Here we used the power iteration algorithm to demonstrate how eigenvectors arise from certain elements of the Jacobian matrix. Examining key Jacobian elements that determine eigenvector structure shows that they originate from a few reaction sensitivities of topologically connected reactions. These reaction sensitivities are at different orders of magnitude, resulting in well-separated dynamics for the metabolites/enzyme forms involved. In a physiologically relevant perturbation, these fast dynamics are not likely to be excited, leaving the slow ones to be main interest of study.

When examining the origin of mode structure, we have introduced a concept we term a half reaction, whose distance from equilibrium is a determinant of the complexity of the mode structure. The half reaction definition arises from linearization of the mass balance equation, where certain reactant/product term has been removed due to differentiation. In a bilinear enzymatic reaction, the reaction sensitivities associated with the substrates/products are often at different orders of magnitude, resulting in half of the reaction responds at a particular time scale while the other half relaxes. This phenomenon is a key feature for the bilinear kinetics occurring in metabolic networks.

## Conclusion

The work here demonstrates an analytical approach to understand kinetic models of metabolism through linear analysis. We showed that diagonal dominance in the Jacobian matrix is an important property in determining simple mode structures and corresponding eigenvalues. We showed the origin of diagonal dominance in terms of specific kinetic parameters of the model. We also described how more complicated mode structures are determined by topologically connected Jacobian elements of similar magnitude. We demonstrated that complicated mode structure arises from the fact that half reaction is close to equilibrium and the presence of connected reaction sensitivities of similar magnitude in **G**. With the recognition that the Jacobian matrix can be factored into **S** and **G**, explicitly representing the chemical, thermodynamic and kinetic properties of a network respectively (30), we can now seek a fundamental understanding of the formation of time scale hierarchies. Such hierarchies can be translated into subspaces of these matrices that in turn give the underpinnings for time-scale decomposition of network functions. Here we demonstrate the existence of such relationships, but a more rigorous mathematical treatment needs to be developed.

## Author Contributions

BOP, BD and DCZ conceived and designed the study. BD and DCZ performed the research and analyzed the data. BD, DCZ and BOP wrote the paper. All authors read and approved the final content.

## Acknowledgments

This work was supported by the Novo Nordisk Foundation (NNF16CC0021858).

